# Serial assessment of iron in the motor cortex in limb-onset Amyotrophic Lateral Sclerosis using Quantitative Susceptibility Mapping

**DOI:** 10.1101/865709

**Authors:** Anjan Bhattarai, Zhaolin Chen, Phillip G. D. Ward, Paul Talman, Susan Mathers, Thanh G Phan, Caron Chapman, James Howe, Sarah Lee, Yennie Lie, Gary F Egan, Phyllis Chua

**Author notes:** Correspondence to: Anjan Bhattarai; Monash Biomedical Imaging, Monash University, 770 Blackburn Rd, Clayton, VIC 3168, Australia.

## Abstract

**Objective:** Dysregulation of iron in the cerebral motor areas has been hypothesized to occur in individuals with Amyotrophic Lateral Sclerosis (ALS). There is still limited knowledge regarding iron dysregulation in the progression of ALS pathology. Our objectives were to use magnetic resonance based Quantitative Susceptibility Mapping (QSM) to investigate the association between iron dysregulation in the motor cortex and clinical manifestations in patients with limb-onset ALS, and to examine changes in the iron concentration in the motor cortex in these patients over a six-month period.

**Methods:** Iron concentration was investigated using magnetic resonance based -QSM in the primary motor cortex and the pre-motor area in thirteen limb-onset ALS patients (including five lumbar onset, six cervical onset and two flail arm patients), and eleven age and sex-matched healthy controls. Nine ALS patients underwent follow-up scans at six months.

**Results:** Significantly increased QSM was observed in the left posterior primary motor area (p = 0.02, Cohen’s d = 0.9) and right anterior primary motor area (p = 0.02, Cohen’s d = 0.92) in all individuals with limb-onset ALS compared to healthy controls. Increased QSM was observed in the primary motor and pre-motor area at baseline in patients with lumbar onset ALS patients, but not cervical limb-onset ALS patients, compared to healthy controls. No significant change in QSM was observed at the six-month follow-up scans in the ALS patients.

**Conclusions:** The findings suggest that iron dysregulation can be detected in the motor cortex in limb-onset ALS, which does not appreciably change over a further 6 months. Individuals with lumbar onset ALS appear to be more susceptible to motor cortex iron dysregulation compared to the individuals with cervical onset ALS. Importantly, this study highlights the potential use of QSM as a radiological indicator in disease diagnosis, and in clinical trials in limb-onset ALS and its subtypes.

**Highlights:**

- Serial measurement of QSM in the motor cortex in limb-onset ALS was performed
- QSM changes in the motor cortex in ALS sub-groups were investigated
- Higher QSM was observed in the motor cortex in Lumbar ALS relative to controls
- QSM is sensitive to iron dysregulation in the motor cortex in limb-onset ALS

## 1. Introduction

Amyotrophic Lateral Sclerosis (ALS) is an idiopathic, fatal and relentlessly progressive neurodegenerative disease which is known to affect upper (UMN) and lower (LMN) motor neurons (Kiernan et al., 2011). Corticospinal tract (CST) degeneration and grey matter neuronal degeneration of the spinal cord, brain stem and motor cortex are considered to be the classical neuropathological features of ALS (Wang, Melhem, Poptani, & Woo, 2011). The pathogenesis is largely unknown. A range of studies indicate that ALS neurodegeneration results from a complex interplay of excitotoxicity, oxidative stress, neuroinflammation, dysfunction of critical proteins and multiple genetic factors (Bonafede & Mariotti, 2017; Carri, Ferri, Cozzolino, Calabrese, & Rotilio, 2003; S. Wang et al., 2011).

A number of studies have suggested iron homeostasis is impaired in the ALS brain (Hadzhieva et al., 2013; Oshiro, Morioka, & Kikuchi, 2011). An *ex vivo* study reported increased iron accumulation, a prominent hallmark of neuroinflammation (Nnah & Wessling-Resnick, 2018) in the microglia of the motor cortex in ALS patients (Kwan et al., 2012). Increased level of motor cortex iron accumulation has been associated with oxidative damage via the Fenton reaction (Contestabile, 2011; Jomova, Vondrakova, Lawson, & Valko, 2010). The Fenton reaction results in the formation of reactive oxygen species (hydroxyl radical) from a weak oxidant (hydrogen peroxide) in the presence of iron (ferrous ion) (Contestabile, 2011). The eventual result is neuronal cell degeneration (Petri, Korner, & Kiaei, 2012). Further support for a deleterious effect of elevated iron in ALS has been found with a therapeutic effect of iron chelation in the superoxide dismutase-1 (SOD-1) mouse model of ALS (Q. Wang et al., 2011). Given the evidence which suggests iron may play a significant role in the pathological process of ALS, an *in vivo* assessment of iron in disease progression in individuals with ALS may provide a pharmacodynamic marker for clinical trials.

Magnetic Resonance Imaging (MRI) has the capability to generate image contrast based on tissue magnetization that can map the neuroanatomical distribution of iron in the human brain. The development of Quantitative Susceptibility Mapping (QSM) has enabled investigations of brain iron deposition *in vivo* (Marques & Bowtell, 2005; Salomir, de Senneville, & Moonen, 2003). QSM is a relatively new MR neuroimaging technique that allows the quantification of magnetic susceptibility using the phase of the MR signal (Haacke et al., 2015). The magnetic susceptibility of tissue perturbs the apparent strength of an externally applied magnetic field and produce a detectable change in the phase of the MRI signal. Acquisition and post-processing MRI techniques have been developed to mathematically transform the non-local phase signal changes into voxel-based estimates of magnetic susceptibility in the brain. The QSM values are linearly proportionate to the concentration of paramagnetic and diamagnetic species in a voxel (Duyn & Schenck, 2017), measured in parts per million (ppm) or parts per billion (ppb). The deposition of paramagnetic (or diamagnetic) species results in increased (or decreased) signal intensity in the QSM image. Iron within brain tissue is predominantly found in the paramagnetic form, and through QSM imaging has been quantified in a number of neurodegenerative diseases (Duyn & Schenck, 2017).

To date there have been seven published cross-sectional studies which have investigated iron in ALS using QSM alone and in combination with other MRI modalities (Acosta-Cabronero et al., 2018; Costagli et al., 2016; Donatelli et al., 2019; Lee et al., 2017; Schweitzer et al., 2015; Weidman et al., 2018; Welton et al., 2019). However, no longitudinal study has been undertaken to investigate changes in QSM values in ALS patients. Longitudinal QSM studies are needed to examine the role of iron in disease progression in ALS and to investigate potential differences in disease progression across the clinical ALS phenotypes.

This study aimed to determine the changes in iron concentration in the motor cortex of limb-onset ALS patients in two clinical phenotypes; namely lumbar onset ALS and cervical onset ALS. In addition, we aimed to determine whether changes in iron concentration over 6 months in the motor cortex were associated with clinical progression in ALS. We hypothesized that iron concentration in the motor cortex in limb-onset ALS would be increased compared to healthy controls, and secondly that iron accumulation in the motor cortex in ALS patients would increase as the symptoms progressed. Furthermore, we hypothesized that there would be measurable changes in the concentration of iron in the motor cortex in ALS patients after six months of disease progression.

## 2. Materials and methods

Ethics approval for this study was obtained from Calvary Health Care Bethlehem and Monash University Research Ethics Committees.

### 2.1. Participants

We recruited participants over 18 years of age, with limb onset ALS who were able to give informed consent themselves. Thirteen (2 females, 11 males) participants with limb onset ALS (cervical onset, n=6, lumbar onset n=5, flail arm onset=2) and eleven (2 females, 9 males) age and gender matched healthy controls underwent MRI scans. Nine patients with ALS participated in follow-up MR scans that were acquired six months after the initial scans. MR scans were not able to be successfully acquired for the other four ALS participants for the following reasons: one participant could not lie flat inside the MRI scanner for the follow-up scan, and data for the other three participants was irreversible corrupted by a hardware storage failure. The participants with ALS underwent neurological assessments with the following clinical measures collected: the laterality and limb of onset, disease duration, UMN and LMN involvement and ALSFRS-R as a measure of disease severity.

### 2.2. MRI acquisition

MRI data for the baseline and follow-up scans were acquired using a 3-Tesla Skyra MRI (Siemens, Erlangen, Germany) equipped with a 32-channel head-and-neck coil. The protocol included a T1-weighted magnetization prepared rapid gradient echo (MPRAGE) and a T2*-weighted gradient-recalled echo (GRE). The MPRAGE acquisition was performed using the following parameters: acquisition time = 5:20 minutes, repetition time = 2300ms, echo time = 2.07ms, flip angle = 9°, field-of-view = 256mm, voxel size = 1 × 1 × 1 mm^3^, 192 slices per volume. The GRE sequence was performed with the following parameters: acquisition time = 5.27 minutes, repetition time = 30ms, echo time = 20ms, flip angle = 15°, field-of-view = 230mm, voxel size = 0.9 × 0.9 × 1.8 mm^3^, 72 slices per volume.

### 2.3. MRI analyses

#### 2.3.1. QSM processing

The phase and QSM processing was carried out using STI-Suite v2.2 (https://people.eecs.berkeley.edu/~chunlei.liu/software.html). Phase and magnitude images from each channel (coil) were reconstructed offline from raw GRE k-space data. T2*-weighted magnitude images were calculated using a sum-of-squares coil combination for registration purposes. Bias correction was performed using N4 from Advanced Normalisation Tools (ANTs) (Tustison et al., 2010). Brain masks were obtained using FSL-BET (Smith, 2002). The phase images from each coil were processed to remove phase warps using Laplacian Unwarping (voxel padding 12×12×12) (Li, Wu, & Liu, 2011; Schofield & Zhu, 2003). Background field was removed using V-SHARP (Schweser, Deistung, Lehr, & Reichenbach, 2011; Wu, Li, Guidon, & Liu, 2012). The processed phase images were combined using a magnitude-weighted average to produce a single-phase image. QSM calculation was then performed using the improved sparse linear equation and least-squares algorithm (iLSQR) (Li et al., 2015). In order to remove air tissue boundary artefacts in QSM, the brain mask was eroded using a spherical kernel with 5mm radius and multiplied with QSM image. QSM values were referenced to the whole brain mean magnetic susceptibility (Straub et al., 2017), and quantified as the mean in each Region of Interest (ROI). In order to exclude the susceptibility contributions from non-grey matter tissues, the mean QSM in each ROI was calculated including the voxels with magnetic susceptibility greater than 10ppb (Schweitzer et al., 2015).

#### 2.3.2. Region of interests (ROIs)

Subject-specific Brodmann area (BA) labels were obtained from the cortical parcellation of anatomical T1-weighted image using FreeSurfer (v6.0.0) (Fischl, 2012; Fischl et al., 2008). Two regions of interest (ROI) were used in the analysis: BA 4 (primary motor) and BA 6 (pre-motor). FreeSurfer cortical parcellation was performed on MASSIVE(Goscinski et al., 2014) using the “recon-all” function. The ROIs were: left and right anterior primary motor cortex (BA 4a), left and right anterior primary motor cortex (BA 4p) and pre-motor motor area (BA 6). The individual ROIs were checked and verified by an experienced neurologist (TP).

#### 2.3.3. Image registration

For the registration of the ROIs to the QSM space, each participant’s T1-weighted image was linearly co-registered to their T2*-weighted magnitude image using ANTs (Avants, Epstein, Grossman, & Gee, 2008). The transformation matrices obtained from the linear registration were then applied to the motor cortex ROIs, originally in their respective MPRAGE space, to bring them to the participant’s QSM space. For visual inspection and comparison, QSM images were registered to standard Montreal Neurological Institute (MNI) space using ANTs (Avants et al., 2008) (Figure 2).

**Figure 1:**
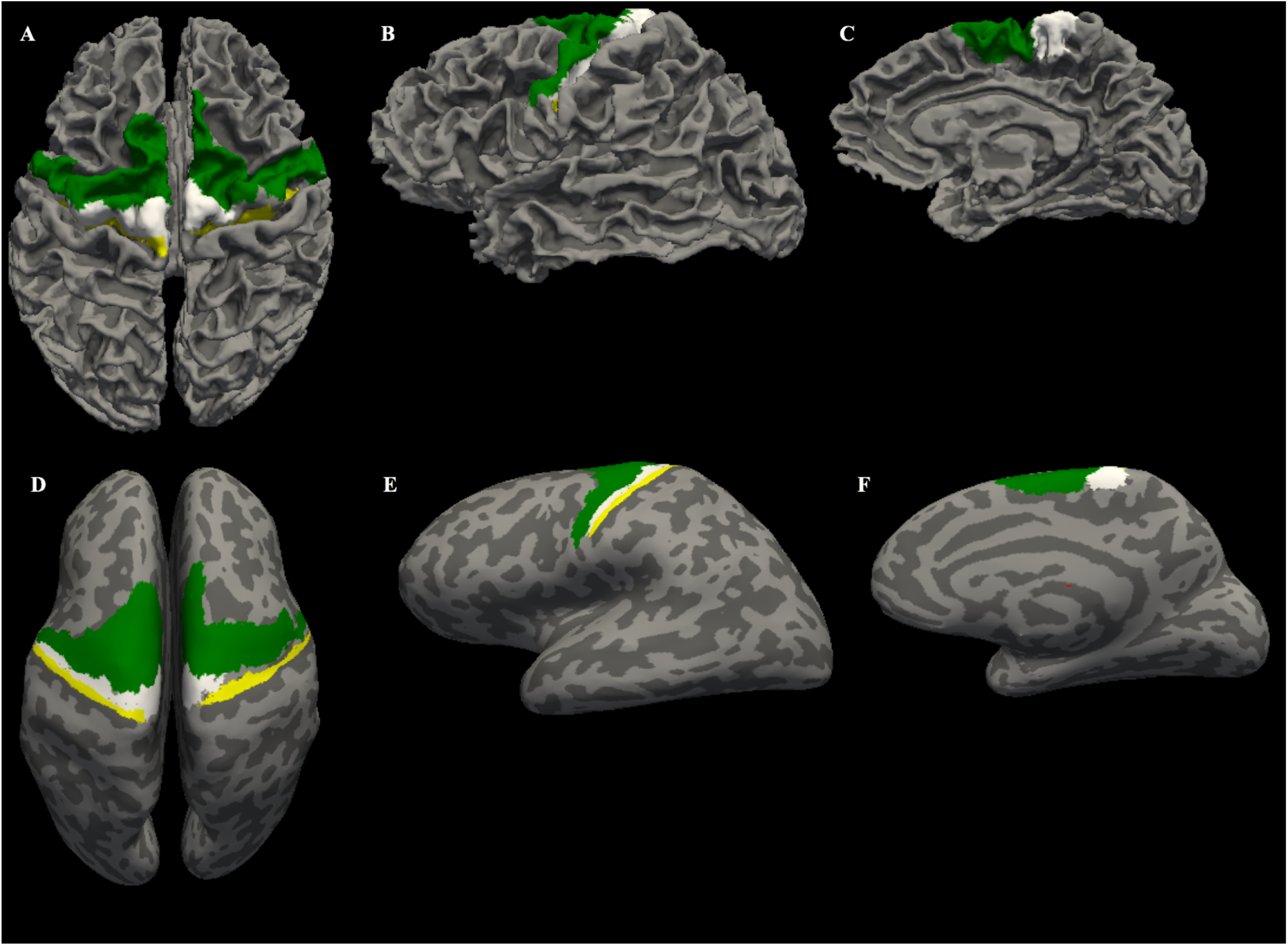
T1-weighted image based automated parcellation of the primary motor area (BA 4) and pre-motor area (Brodmann area 6) using FreeSurfer. The labels are anterior BA 4 (white), posterior BA 4 (yellow), pre-motor area (green). A, B and C shows top view, lateral view, and midsagittal view respectively of normal surface. Panels D, E and F show top view, lateral view, and midsagittal view respectively of the inflated surface.

**Figure 2:**
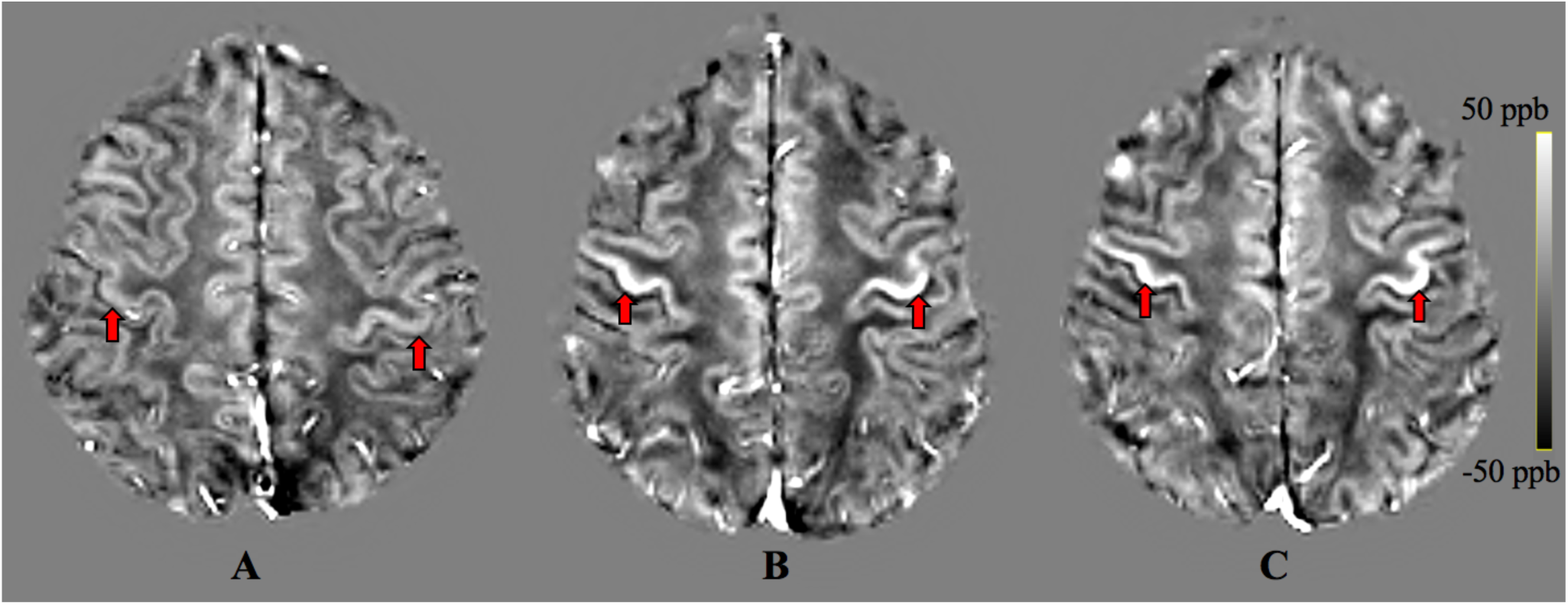
Hyperintense signal in the primary motor cortex (red arrows) in group-average QSM images for (A) a healthy control, (B) an ALS patient at study entry, and (C) the same ALS patient at the 6 months follow-up scan.

#### 2.3.4. Statistical Analyses

The ALS patients at baseline were further categorised into two groups on the basis of their individual clinical phenotype as either lumbar onset ALS or cervical onset ALS. Due to the small number of ALS patients with a follow-up scan, the patients with six-month follow-up scans were not categorised on the basis of clinical phenotype and were analysed as a single group. Age related effects in the QSM measures were estimated in the control participant cohort and used to correct for normal age-related changes in the ALS patient cohorts. Group differences in QSM values between the ALS clinical phenotype cohorts and the healthy control participants were examined in the primary motor and pre-motor area ROIs. To test whether QSM was increased in all limb onset ALS patients (lumbar onset, cervical onset, flail arm ALS) compared to the heathy controls a non-parametric Wilcoxon rank sum test was used with the reported p-values calculated in MATLAB using the “ranksum” function. The p-value significance level was p<0.05 using a one-tailed distribution, and the comparisons between the ALS subgroups (lumbar and cervical onset ALS) with healthy controls were corrected for multiple comparisons. The one-tailed distribution was chosen based an *ex vivo* study (Kwan et al., 2012) and other QSM studies, (Costagli et al., 2016; Lee et al., 2017) which have found iron concentration to be elevated in the primary motor cortex of ALS patients compared to healthy controls. To test whether QSM was changed at six-month follow-up compared to baseline, a non-parametric Wilcoxon signed rank test was used with the reported p values (two-tailed) calculated in MATLAB using the “signrank” function. The p-value significance level was p<0.05. The sample is paired such that QSM observations at baseline were considered only from the individuals whose follow-up observations were available.

Adjusted p-values were calculated in MATLAB controlling the false discovery rate (FDR) following the Benjamini & Hochberg (1995) procedure (Benjamini & Hochberg, 1995). Unadjusted p-values were reported in the following two instances. First, when comparing the baseline QSM values in the ALS group (lumbar onset, cervical onset, flail arm ALS) with healthy controls. Second, when comparing the baseline QSM values and at six-month follow-up due to the exploratory nature of the analysis with the small number of ALS patients. The QSM values are reported as group mean ± standard error of mean (SEM). The effect size was calculated using Cohen’s d (Cohen, 2013). The assumption of paired sample was considered to calculate the effect sizes while comparing baseline and follow-up QSM observations.

Correlations with disease duration and disease severity were assessed using non-parametric Spearman’s rho (two-tailed). Disease duration was defined as the time between the date of disease onset based on patient report of symptom onset and the date of the MRI scan. Disease severity was quantified using (ALSFRS-R) scale (Cedarbaum et al., 1999). ALSFRS-R scores range from 0 to 48, where a score of 0 represents severely impaired and a score of 48 is unimpaired. All other parameters used in statistical analyses were set to default unless specified in the text. This study is underpowered to draw an inference from the correlation analyses with clinical measures but statistics are reported in the supplement to facilitate future meta-analyses of this rare condition and its clinical subtypes.

## 3. Results

Significantly increased QSM values were observed in individuals with limb-onset ALS compared to healthy controls in the left posterior primary motor cortex and right anterior primary motor cortex (Table 2). Increased QSM values were observed in all other motor cortex regions of limb-onset ALS participants compared to healthy controls but did not reach the level of statistical significance.

**Table 1:** Participant demographics and clinical scores (mean ± standard deviation)

|  | ALS | Controls |
| --- | --- | --- |
| Gender (F:M) | 2:11 | 2:9 |
| Age at baseline (years) | 56.2 ± 12.6 | 52.3 ± 11.6 |
| Age at follow-up (years) | 59.5 ± 11.0 | NA |
| Number of participants at baseline | 13 | 11 |
| Number of participants at follow-up | 13 (data available from 9) | NA |
| Onset (Cervical: Lumbar: Flail Arm) | 6:5:2 | NA |
| Disease duration at baseline (years)<sup>1</sup> | 3.8 ± 3.8 | NA |
| ALSFRS-R1 (scores available from 11 participants) | 37.8 ± 6.2 | NA |
| ALSFRS-R2 (scores available from 8 participants) | 34.8 ± 9.4 | NA |

<sup>1</sup>-Disease duration based on patient report of symptom onset to time of baseline scan
<sup>2</sup>-ALSFRS-R1, and ALSFRS-R2 are ALSFRS-R scores at baseline and 6 months follow-up respectively.

**Table 2:** Summary of the cross-sectional age-corrected QSM observations. Statistical significance measured with Wilcoxon rank sum test (1-tailed).

| ROI | QSM (ppb) |  | Baseline ALS vs Health Controls |  |
| --- | --- | --- | --- | --- |
| | Healthy Control mean ± SEM | Baseline ALS mean ± SEM | p value | Cohen's d |
| Left primary Motor Area (Anterior) | 27.8 ± 0.9 | 29.6 ± 1.0 | 0.12 | 0.57 |
| Right Primary Motor Area (Anterior) | 29.7 ± 1.1 | 33.0 ± 0.9 | 0.02* | 0.92 |
| Left Primary Motor Area (Posterior) | 27.0 ± 0.8 | 30.2 ± 1.2 | 0.02* | 0.9 |
| Right Primary Motor Area (Posterior) | 28.3 ± 0.8 | 30.3 ± 1.1 | 0.09 | 0.58 |
| Left Pre-Motor Area | 23.1 ± 0.6 | 24.4 ± 0.8 | 0.23 | 0.50 |
| Right Pre-Motor Area | 23.2 ± 0.5 | 24.6 ± 1.0 | 0.09 | 0.47 |

We observed an increased pattern of QSM values in the anterior primary motor area (bilateral), left posterior primary motor area and the right pre-motor area at baseline in patients with lumbar onset ALS compared to healthy controls (Table 3) together with strong effect sizes (Cohen’s d). The group differences in QSM values between the cervical onset ALS patients at baseline and the healthy controls were not statistically significant (Table 3). The change in QSM values in limb-onset ALS participants between baseline and six-month follow-up scans was not significant. (Table 4).

**Table 3:** Summary of the cross-sectional age-corrected QSM observations between healthy controls and lumbar and cervical onset ALS respectively at baseline. Statistical significance measured with Wilcoxon rank sum test (1-tailed).

| ROI | QSM (ppb) |  |  | Lumbar onset ALS Vs Healthy Controls |  | Cervical onset ALS vs Healthy Controls |  |
| --- | --- | --- | --- | --- | --- | --- | --- |
| | Healthy Control Mean ± SEM | Lumbar onset ALS Mean ± SEM | Cervical onset ALS Mean ± SEM | p value adjusted | Cohen's d | p value adjusted | Cohen's d |
| Left primary Motor Area (Anterior) | 27.8 ± 0.9 | 30.8 ± 1.1 | 29.2 ± 1.8 | 0.07 | 1.09 | 0.4 | 0.4 |
| Right Primary Motor Area (Anterior) | 29.7 ± 1.1 | 34.4 ± 1.9 | 32.5 ± 1.2 | 0.07 | 1.21 | 0.36 | 0.81 |
| Left Primary Motor Area (Posterior) | 27.0 ± 0.8 | 30.5 ± 1.32 | 29.8 ± 2.5 | 0.07 | 1.34 | 0.4 | 0.70 |
| Right Primary Motor Area (Posterior) | 28.3 ± 0.8 | 31.2 ± 1.5 | 30.2 ± 2.0 | 0.07 | 1.01 | 0.4 | 0.52 |
| Left Pre-Motor Area | 23.1 ± 0.6 | 25.9 ± 2.0 | 23.5 ± 0.6 | 0.13 | 0.94 | 0.4 | 0.22 |
| Right Pre-Motor Area | 23.2 ± 0.53 | 26.6 ± 2.2 | 23.1 ± 0.6 | 0.07 | 1.12 | 0.4 | -0.08 |

**Table 4:** Summary of the serial age-corrected QSM observations. Statistical significance measured with Wilcoxon signed rank test (2-tailed). The assumption of paired sample was considered while calculating the effect sizes.

| ROI | QSM(ppb) |  | Baseline ALS vs 6 months |  |
| --- | --- | --- | --- | --- |
| | Baseline ALS mean ± SEM | 6 months ALS mean ± SEM | p value | Cohen's d |
| Left primary Motor Area (Anterior) | 30 ± 1.3 | 29.9 ± 1.3 | 1 | -0.03 |
| Right Primary Motor Area (Anterior) | 33.7 ± 1.2 | 34.4 ± 1.9 | 0.73 | 0.23 |
| Left Primary Motor Area (Posterior) | 31.2 ± 1.6 | 29.9 ± 1.4 | 0.05 | -0.77 |
| Right Primary Motor Area (Posterior) | 31 ± 1.5 | 31.3 ± 2 | 0.91 | 0.13 |
| Left Pre-Motor Area | 24.6 ± 1.2 | 25.2 ± 1.3 | 0.3 | 0.56 |
| Right Pre-Motor Area | 25.1 ± 1.4 | 25.7 ± 1.4 | 0.05 | 0.76 |

**Table 5:** Summary of the correlations between clinical measures (disease duration and disease severity i.e., ALSFRS-R score) and age corrected QSM observations.

| ROI | Disease duration vs QSM |  |  |  |  |  |  |  | ALSFRS_R vs QSM |  |  |  |  |  |  |  |
| --- | --- | --- | --- | --- | --- | --- | --- | --- | --- | --- | --- | --- | --- | --- | --- | --- |
|  | Correlation at baseline |  |  |  |  |  | Correlation at follow-up |  | Correlation at baseline |  |  |  |  |  | Correlation at follow-up |  |
|  | ALS |  | Lumbar Onset ALS |  | Cervical Onset ALS |  | ALS |  | ALS |  | Lumbar Onset ALS |  | Cervical Onset ALS |  | ALS |  |
|  | rho | p | rho | p | rho | p | rho | p | rho | p | rho | p | rho | p | rho | p |
| Left primary Motor Area (Anterior) | 0.09 | 0.76 | 0 | 1 | 0.54 | 0.3 | 0.4 | 0.29 | 0.43 | 0.18 | 0.8 | 0.33 | -0.14 | 0.8 | -0.05 | 0.92 |
| Right Primary Motor Area (Anterior) | 0.08 | 0.81 | -0.1 | 0.95 | 0.31 | 0.56 | 0.23 | 0.55 | 0.37 | 0.26 | 0.8 | 0.33 | 0.09 | 0.92 | -0.01 | 0.99 |
| Left Primary Motor Area (Posterior) | 0.27 | 0.37 | 0.7 | 0.23 | -0.03 | 1 | 0.2 | 0.61 | 0.39 | 0.24 | 1 | 0.08 | 0.09 | 0.92 | -0.13 | 0.76 |
| Right Primary Motor Area (Posterior) | 0.15 | 0.63 | -0.1 | 0.95 | 0.14 | 0.8 | 0.23 | 0.55 | 0.38 | 0.25 | 0.6 | 0.42 | 0.31 | 0.56 | 0.08 | 0.85 |
| Left Pre-Motor Area | 0.03 | 0.94 | -0.1 | 0.95 | 0.26 | 0.66 | 0.13 | 0.74 | 0.32 | 0.34 | 0.8 | 0.33 | -0.43 | 0.42 | 0.02 | 0.96 |
| Right Pre-Motor Area | 0.15 | 0.62 | -0.2 | 0.78 | 0.6 | 0.24 | 0.42 | 0.27 | 0.66 | 0.03* | 0.6 | 0.42 | 0.49 | 0.36 | 0.55 | 0.16 |

Positive correlations were observed between disease duration and QSM values in all ROIs in limb-onset ALS at both baseline and follow-up but were not statistically significant (statistics are reported in supplement). Correlations between disease duration at baseline and QSM observations in lumbar and cervical onset ALS at baseline were not statistically significant. Progression in clinical symptoms did not reflect in the change in QSM observations as we did not observe statistically significant anti-correlation between disease severity scores as measured by ALSFRS-R scale and QSM observations in limb-onset ALS and its subtypes. The statistics for correlation analyses are reported in the supplement.

## 4. Discussion

To the best of our knowledge, this is the first study to investigate motor cortex iron dysregulation in limb-onset ALS using MR-based QSM and its two clinical subtypes: lumbar onset ALS and cervical onset ALS. Our findings suggest that increased iron level can be detected significantly in the motor cortex in limb-onset ALS relative to healthy controls. We observed that individuals with lumbar onset ALS had more evidence of motor cortex iron dysregulation than cervical onset ALS relative to healthy controls. The results suggest that QSM is sensitive to ALS subtypes, and may aid as a radiological marker in clinical trials and disease diagnosis. We observed no detectable change in the iron concentration over 6 months of disease, and no significant correlation with disease duration. This may suggest that motor cortex iron dysregulation is present in the earlier stage and does not progress significantly over the course of disease in limb-onset ALS. It also provides guidance to its use as a marker of disease progression in interventional studies as to minimum observation periods (>6 months) to expect changes.

Although limited with small sample size, this is the first study to investigate serial assessment of motor cortex iron dysregulation in ALS. Given the heterogeneous presentation of ALS, our cohort was restricted to limb-onset ALS to enable a more focused exploration of the most common ALS clinical phenotype. Limb-onset ALS also has a longer duration of illness (Turner et al., 2010; Turner, Parton, Shaw, Leigh, & Al-Chalabi, 2003) such that the majority of patients were able to be rescanned at 6 months.

Iron plays a significant role as a cofactor in many regulatory enzymes involved in the cellular metabolism process occurring in the mitochondria (Kwan et al., 2012). Due to the high metabolic activity of the brain, iron has a prominent role in the central nervous system. Iron levels in the brain have been reported to increase as part of the normal ageing process (Zecca, Youdim, Riederer, Connor, & Crichton, 2004). However, increased iron levels have also been observed in people diagnosed with neurodegenerative disorders such as Parkinson’s disease (Zecca et al., 2004), Friedreich Ataxia (Ward et al., 2019) and Alzheimer’s disease (Rouault & Cooperman, 2006). A pathological study conducted in two ALS patients (42 and 51 years) reported increased iron accumulation in the motor cortex despite their young age (Kwan et al., 2012). Imaging findings of the same study showed hypointense signal in the deep layers of motor cortex in 7 Tesla MRI T2*-weighted gradient echo images in both patients, suggesting that the MRI signal alterations were due to the increased iron accumulation. Our study corroborates the findings of a cross-sectional study by Lee et al. (2017) which reported significantly higher QSM in the motor cortex of ALS patients compared to healthy controls (Lee et al., 2017). Another cross-sectional study by Costagli et al. (2016) reported significantly higher magnetic susceptibility in M1 subregions of ALS patients compared to healthy controls (Costagli et al., 2016). They demonstrated a strong positive correlation between magnetic susceptibility and the expected concentration of iron in different cortical regions in healthy controls. The expected concentration of iron in healthy controls was calculated based on the findings of an earlier histological study (Hallgren & Sourander, 1958). Their results were consistent with previous *ex vivo* findings which reported increased iron accumulation in microglial cells in the deep layers of the motor cortex (Kwan et al., 2012). Qualitatively, the diagnostic accuracy of QSM in ALS compared to earlier MRI techniques such as T2-weighted, T2*-weighted and T2 FLAIR images has been demonstrated by Schweitzer et al. (2015), showing significantly greater motor cortex susceptibility in motor neurone disease (MND) patients (including both ALS and primary lateral sclerosis) which was postulated to reflect increased iron accumulation (Schweitzer et al., 2015). Primary lateral sclerosis (PLS) is a less common progressive neurodegenerative disease which is known to affect upper motor neurons (Tartaglia et al., 2007). Our study results are consistent with these studies which indicate iron dysfunction in the motor cortex in ALS. Furthermore, the difference noted in lumbar onset ALS highlights the importance of studying subgroups of ALS as the underlying iron mediated pathological mechanisms may be different.

There are methodological considerations to the current study. QSM is a nascent method undergoing rapid development (Langkammer et al., 2018) and importantly QSM provides relative rather than absolute values of magnetic susceptibility measurements. The k-space origin of the calculated susceptibility map during the QSM reconstruction process is mathematically unconstrained. As a result, the QSM susceptibility map has an unknown offset so the susceptibility values must be referenced to a particular brain region or tissue (Deistung, Schweser, & Reichenbach, 2017). In this study, the QSM values reported were referenced to whole brain mean susceptibility value rather than the susceptibility of specific anatomical region, in order to minimize any potential biases that may result from disease related susceptibility changes in a specific reference region. However, it is unknown whether the whole brain mean appreciably changes longitudinally in ALS and any such change may obscure the findings of this study. The QSM values between studies may also differ depending upon the methods and parameters used for phase processing, as well as the magnetic field strength, imaging protocols and equipment used for MRI data acquisition. These factors can impact on the QSM values between different published studies but do not have any impact on the significance of group difference and longitudinal changes within a single study. Further developments towards protocol and processing standardization may mitigate these factors and improve the reliability of QSM analyses from multi-center studies.

The current study has several limitations. The study was conducted in relatively small sized cohorts of patients with clinical possible, probable or definite limb-onset ALS. Follow-up MRI data was available for only nine out of thirteen patients. Whilst larger sample sizes would contribute to higher statistical power for between group analyses, the rarity and rapid progression of the disease make large cohort studies difficult. For the longitudinal analysis, MRI data was available for two time points with a relatively short duration of six-month between the scans. Although we chose to study only limb-onset ALS, the cohort was still heterogeneous with respect to disease stages disease duration and clinical symptoms of ALS. Future longitudinal studies over a longer time span are needed to examine the trajectory of iron accumulation in ALS, although unfortunately such studies will be limited by the short prognosis in most patients with ALS and the concomitant difficulty of lying flat in the MRI scanner as the disease progresses.

## 5. Conclusion

This study demonstrates the efficacy of QSM to detect iron concentration differences in the motor cortex of individuals with limb-onset ALS. The results suggest that iron levels are significantly increased in the primary motor cortex in limb-onset ALS relative to healthy controls, and the changes may not show significant progression over a six-month period. Increased iron concentration was also pronounced in the pre-motor areas in ALS patients compared to healthy controls. Individuals with lumbar onset ALS had more evidence of motor cortex iron dysregulation than individuals with cervical onset relative to healthy controls. Analysis of iron concentration can provide meaningful insight of the pathological process of ALS. Future longitudinal studies with multiple scans, performed over longer period in a larger cohort, grouped on the basis of disease stage, severity, clinical presentations and phenotypes are required to further explore the role of iron in the degeneration process.

## Acknowledgement

We would like to thank all the volunteers who participated in this study. We are also grateful to the radiographers at Monash Biomedical Imaging for their assistance with MRI data collection and our nurse research assistants Ruth Krasniqi and Anna Smith. Funding for this project was obtained through the Monash University Strategic Grant Scheme. AB was supported by The Australian Rotary Health / Rotary Club of Sandy Bay PhD Scholarship in Motor Neuron Disease. We declare no conflict of interest.

## Author Contributions

The first author named is lead and corresponding author. All other authors are listed on the basis of their contribution to the project from most to least with the main investigators last. **Anjan Bhattarai:** Conceptualization, Methodology, Software, Validation, Formal analysis, Investigation, Data curation, Writing – Original Draft, Writing – Review & Editing, Visualization. **Zhaolin Chen:** Conceptualization, Methodology, Validation, Formal analysis, Investigation, Data curation, Writing – Review & Editing, Supervision. **Phillip G.D. Ward:** Methodology, Software, Writing – Review & Editing. **Paul Talman:** Investigation, Resources, Supervision. **Susan Mathers:** Investigation, Resources. **Thanh G Phan:** Validation, Writing – Review & Editing. **Caron Chapman:** Investigation, Resources. **James Howe:** Investigation, Resources. **Sarah Lee:** Investigation, Resources. **Yennie Lie:** Investigation, Resources. **Gary F Egan:** Investigation, Resources, Writing – Review & Editing, Supervision. **Phyllis Chua:** Conceptualization, Methodology, Formal analysis, Investigation, Resources, Data curation, Writing – Review & Editing, Supervision, Project administration.

## Supplementary materials

### Estimation and correction for age related changes in QSM

Age related effects in the QSM measures were estimated in the control participant cohort and used to correct for normal age related changes in the ALS patient cohorts.

**Age corrected QSM = Uncorrected QSM-b × (Age - mean age)**, where b is the slope of a regression and mean age is the mean age of healthy controls.

b is calculated using linear regression in SPSS, choosing uncorrected QSM of healthy controls for each ROI as dependent variable and age of the heathy controls as independent variable. The unstandardized B is the value for b.

### Statistics for clinical correlation

## References

Acosta-Cabronero, J., Machts, J., Schreiber, S., Abdulla, S., Kollewe, K., Petri, S., … Nestor, P. J. (2018). Quantitative Susceptibility MRI to Detect Brain Iron in Amyotrophic Lateral Sclerosis. Radiology, 289(1), 195–203. doi: 10.1148/radiol.2018180112

Avants, B. B., Epstein, C. L., Grossman, M., & Gee, J. C. (2008). Symmetric diffeomorphic image registration with cross-correlation: evaluating automated labeling of elderly and neurodegenerative brain. Med Image Anal, 12(1), 26–41. doi: 10.1016/j.media.2007.06.004

Benjamini, Y., & Hochberg, Y. (1995). Controlling The False Discovery Rate - A Practical And Powerful Approach To Multiple Testing. J. Royal Statist. Soc., Series B, 57, 289–300. doi: 10.2307/2346101

Bonafede, R., & Mariotti, R. (2017). ALS Pathogenesis and Therapeutic Approaches: The Role of Mesenchymal Stem Cells and Extracellular Vesicles. Front Cell Neurosci, 11. doi: 10.3389/fncel.2017.00080

Carri, M. T., Ferri, A., Cozzolino, M., Calabrese, L., & Rotilio, G. (2003). Neurodegeneration in amyotrophic lateral sclerosis: the role of oxidative stress and altered homeostasis of metals. Brain Res Bull, 61(4), 365–374.

Cedarbaum, J. M., Stambler, N., Malta, E., Fuller, C., Hilt, D., Thurmond, B., & Nakanishi, A. (1999). The ALSFRS-R: a revised ALS functional rating scale that incorporates assessments of respiratory function. BDNF ALS Study Group (Phase III). J Neurol Sci, 169(1-2), 13–21. doi: 10.1016/s0022-510x(99)00210-5

Cohen, J. (2013). Statistical Power Analysis for the Behavioral Sciences: Taylor & Francis.

Contestabile, A. (2011). Amyotrophic lateral sclerosis: from research to therapeutic attempts and therapeutic perspectives. Curr Med Chem, 18(36), 5655–5665.

Costagli, M., Donatelli, G., Biagi, L., Caldarazzo Ienco, E., Siciliano, G., Tosetti, M., & Cosottini, M. (2016). Magnetic susceptibility in the deep layers of the primary motor cortex in Amyotrophic Lateral Sclerosis. Neuroimage Clin, 12, 965–969. doi: 10.1016/j.nicl.2016.04.011

Deistung, A., Schweser, F., & Reichenbach, J. R. (2017). Overview of quantitative susceptibility mapping. NMR Biomed, 30(4). doi: 10.1002/nbm.3569

Donatelli, G., Caldarazzo Ienco, E., Costagli, M., Migaleddu, G., Cecchi, P., Siciliano, G., & Cosottini, M. (2019). MRI cortical feature of bulbar impairment in patients with amyotrophic lateral sclerosis. Neuroimage Clin, 24, 101934. doi: 10.1016/j.nicl.2019.101934

Duyn, J. H., & Schenck, J. (2017). Contributions to magnetic susceptibility of brain tissue. NMR Biomed, 30(4). doi: 10.1002/nbm.3546

Fischl, B. (2012). FreeSurfer. Neuroimage, 62(2), 774–781. doi: 10.1016/j.neuroimage.2012.01.021

Fischl, B., Rajendran, N., Busa, E., Augustinack, J., Hinds, O., Yeo, B. T., … Zilles, K. (2008). Cortical folding patterns and predicting cytoarchitecture. Cereb Cortex, 18(8), 1973–1980. doi: 10.1093/cercor/bhm225

Goscinski, W. J., McIntosh, P., Felzmann, U., Maksimenko, A., Hall, C., Gureyev, T., … Egan, G. (2014). The multi-modal Australian ScienceS Imaging and Visualization Environment (MASSIVE) high performance computing infrastructure: applications in neuroscience and neuroinformatics research. Frontiers in Neuroinformatics,8(30). doi: 10.3389/fninf.2014.00030

Haacke, E. M., Liu, S., Buch, S., Zheng, W., Wu, D., & Ye, Y. (2015). Quantitative susceptibility mapping: current status and future directions. Magn Reson Imaging, 33(1), 1–25. doi: 10.1016/j.mri.2014.09.004

Hadzhieva, M., Kirches, E., Wilisch-Neumann, A., Pachow, D., Wallesch, M., Schoenfeld, P., … Mawrin, C. (2013). Dysregulation of iron protein expression in the G93A model of amyotrophic lateral sclerosis. Neuroscience, 230, 94–101. doi: 10.1016/j.neuroscience.2012.11.021

Hallgren, B., & Sourander, P. (1958). The effect of age on the non-haemin iron in the human brain. J Neurochem, 3(1), 41–51. doi: 10.1111/j.1471-4159.1958.tb12607.x

Jomova, K., Vondrakova, D., Lawson, M., & Valko, M. (2010). Metals, oxidative stress and neurodegenerative disorders. Mol Cell Biochem, 345(1-2), 91–104. doi: 10.1007/s11010-010-0563-x

Kiernan, M. C., Vucic, S., Cheah, B. C., Turner, M. R., Eisen, A., Hardiman, O., … Zoing, M. C. (2011). Amyotrophic lateral sclerosis. Lancet, 377(9769), 942–955. doi: 10.1016/s0140-6736(10)61156-7

Kwan, J. Y., Jeong, S. Y., Van Gelderen, P., Deng, H. X., Quezado, M. M., Danielian, L. E., … Floeter, M. K. (2012). Iron accumulation in deep cortical layers accounts for MRI signal abnormalities in ALS: correlating 7 tesla MRI and pathology. PLoS One, 7(4), e35241. doi: 10.1371/journal.pone.0035241

Langkammer, C., Schweser, F., Shmueli, K., Kames, C., Li, X., Guo, L., … Bilgic, B. (2018). Quantitative susceptibility mapping: Report from the 2016 reconstruction challenge. Magn Reson Med, 79(3), 1661–1673. doi: 10.1002/mrm.26830

Lee, J. Y., Lee, Y. J., Park, D. W., Nam, Y., Kim, S. H., Park, J., … Oh, K. W. (2017). Quantitative susceptibility mapping of the motor cortex: a comparison of susceptibility among patients with amyotrophic lateral sclerosis, cerebrovascular disease, and healthy controls. Neuroradiology, 59(12), 1213–1222. doi: 10.1007/s00234-017-1933-9

Li, W., Wang, N., Yu, F., Han, H., Cao, W., Romero, R., … Liu, C. (2015). A method for estimating and removing streaking artifacts in quantitative susceptibility mapping. Neuroimage, 108, 111–122. doi: 10.1016/j.neuroimage.2014.12.043

Li, W., Wu, B., & Liu, C. (2011). Quantitative susceptibility mapping of human brain reflects spatial variation in tissue composition. Neuroimage, 55(4), 1645–1656. doi: 10.1016/j.neuroimage.2010.11.088

Marques, J. P., & Bowtell, R. (2005). Application of a Fourier-based method for rapid calculation of field inhomogeneity due to spatial variation of magnetic susceptibility. Concepts in Magnetic Resonance Part B: Magnetic Resonance Engineering, 25B(1), 65–78. doi: 10.1002/cmr.b.20034

Nnah, I. C., & Wessling-Resnick, M. (2018). Brain Iron Homeostasis: A Focus on Microglial Iron. Pharmaceuticals, 11(4), 129.

Oshiro, S., Morioka, M. S., & Kikuchi, M. (2011). Dysregulation of iron metabolism in Alzheimer’s disease, Parkinson’s disease, and amyotrophic lateral sclerosis. Adv Pharmacol Sci, 2011, 378278. doi: 10.1155/2011/378278

Petri, S., Korner, S., & Kiaei, M. (2012). Nrf2/ARE Signaling Pathway: Key Mediator in Oxidative Stress and Potential Therapeutic Target in ALS. Neurol Res Int, 2012, 878030. doi: 10.1155/2012/878030

Rouault, T. A., & Cooperman, S. (2006). Brain iron metabolism. Semin Pediatr Neurol, 13(3), 142–148. doi: 10.1016/j.spen.2006.08.002

Salomir, R., de Senneville, B. D., & Moonen, C. T. (2003). A fast calculation method for magnetic field inhomogeneity due to an arbitrary distribution of bulk susceptibility. Concepts in Magnetic Resonance Part B: Magnetic Resonance Engineering, 19B(1), 26–34. doi: 10.1002/cmr.b.10083

Schofield, M. A., & Zhu, Y. (2003). Fast phase unwrapping algorithm for interferometric applications. Opt Lett, 28(14), 1194–1196.

Schweitzer, A. D., Liu, T., Gupta, A., Zheng, K., Seedial, S., Shtilbans, A., … Tsiouris, A. J. (2015). Quantitative susceptibility mapping of the motor cortex in amyotrophic lateral sclerosis and primary lateral sclerosis. AJR Am J Roentgenol, 204(5), 1086–1092. doi: 10.2214/ajr.14.13459

Schweser, F., Deistung, A., Lehr, B. W., & Reichenbach, J. R. (2011). Quantitative imaging of intrinsic magnetic tissue properties using MRI signal phase: an approach to in vivo brain iron metabolism? Neuroimage, 54(4), 2789–2807. doi: 10.1016/j.neuroimage.2010.10.070

Smith, S. M. (2002). Fast robust automated brain extraction. Hum Brain Mapp, 17(3), 143–155. doi: 10.1002/hbm.10062

Straub, S., Schneider, T. M., Emmerich, J., Freitag, M. T., Ziener, C. H., Schlemmer, H. P., … Laun, F. B. (2017). Suitable reference tissues for quantitative susceptibility mapping of the brain. Magn Reson Med, 78(1), 204–214. doi: 10.1002/mrm.26369

Tartaglia, M. C., Rowe, A., Findlater, K., Orange, J. B., Grace, G., & Strong, M. J. (2007). Differentiation between primary lateral sclerosis and amyotrophic lateral sclerosis: examination of symptoms and signs at disease onset and during follow-up. Arch Neurol, 64(2), 232–236. doi: 10.1001/archneur.64.2.232

Turner, M. R., Brockington, A., Scaber, J., Hollinger, H., Marsden, R., Shaw, P. J., & Talbot, K. (2010). Pattern of spread and prognosis in lower limb-onset ALS. Amyotroph Lateral Scler, 11(4), 369–373. doi: 10.3109/17482960903420140

Turner, M. R., Parton, M. J., Shaw, C. E., Leigh, P. N., & Al-Chalabi, A. (2003). Prolonged survival in motor neuron disease: a descriptive study of the King’s database 1990-2002. J Neurol Neurosurg Psychiatry, 74(7), 995–997. doi: 10.1136/jnnp.74.7.995

Tustison, N. J., Avants, B. B., Cook, P. A., Zheng, Y., Egan, A., Yushkevich, P. A., & Gee, J. C. (2010). N4ITK: improved N3 bias correction. IEEE Trans Med Imaging, 29(6), 1310–1320. doi: 10.1109/tmi.2010.2046908

Wang, Q., Zhang, X., Chen, S., Zhang, X., Zhang, S., Youdium, M., & Le, W. (2011). Prevention of motor neuron degeneration by novel iron chelators in SOD1(G93A) transgenic mice of amyotrophic lateral sclerosis. Neurodegener Dis, 8(5), 310–321. doi: 10.1159/000323469

Wang, S., Melhem, E. R., Poptani, H., & Woo, J. H. (2011). Neuroimaging in amyotrophic lateral sclerosis. Neurotherapeutics, 8(1), 63–71. doi: 10.1007/s13311-010-0011-3

Ward, P. G. D., Harding, I. H., Close, T. G., Corben, L. A., Delatycki, M. B., Storey, E., … Egan, G. F. (2019). Longitudinal evaluation of iron concentration and atrophy in the dentate nuclei in friedreich ataxia. Mov Disord, 34(3), 335–343. doi: 10.1002/mds.27606

Weidman, E. K., Schweitzer, A. D., Niogi, S. N., Brady, E. J., Starikov, A., Askin, G., … Tsiouris, A. J. (2018). Diffusion tensor imaging and quantitative susceptibility mapping as diagnostic tools for motor neuron disorders. Clin Imaging, 53, 6–11. doi: 10.1016/j.clinimag.2018.09.015

Welton, T., Maller, J. J., Lebel, R. M., Tan, E. T., Rowe, D. B., & Grieve, S. M. (2019). Diffusion kurtosis and quantitative susceptibility mapping MRI are sensitive to structural abnormalities in amyotrophic lateral sclerosis. Neuroimage Clin, 24, 101953. doi: 10.1016/j.nicl.2019.101953

Wu, B., Li, W., Guidon, A., & Liu, C. (2012). Whole brain susceptibility mapping using compressed sensing. Magn Reson Med, 67(1), 137–147. doi: 10.1002/mrm.23000

Zecca, L., Youdim, M. B., Riederer, P., Connor, J. R., & Crichton, R. R. (2004). Iron, brain ageing and neurodegenerative disorders. Nat Rev Neurosci, 5(11), 863–873. doi: 10.1038/nrn1537

